# Male sex hormones increase excitatory neuron production in developing human neocortex

**DOI:** 10.1101/2020.10.24.353359

**Authors:** Iva Kelava, Ilaria Chiaradia, Laura Pellegrini, Alex T. Kalinka, Madeline A. Lancaster

## Abstract

The presence of male-female brain differences has long been a controversial topic. Yet simply negating the existence of biological differences has detrimental consequences for all sexes and genders, particularly for the development of accurate diagnostic tools, effective drugs and understanding of disease. The most well-established morphological difference is size, with males having on average a larger brain than females; yet a mechanistic understanding of how this difference arises remains to be elucidated. Here, we use brain organoids to test the roles of sex chromosomes and sex steroids during development. While we show no observable differences between XX and XY brain organoids, sex steroids, namely androgens, increase proliferation of cortical neural progenitors. Transcriptomic analysis reveals effects on chromatin remodelling and HDAC activity, both of which are also implicated in the male-biased conditions autism spectrum disorder and schizophrenia. Finally, we show that higher numbers of progenitors result specifically in increased upper-layer excitatory neurons. These findings uncover a hitherto unknown role for male sex hormones in regulating excitatory neuron number within the human neocortex and represent a first step towards understanding the origin of human sex-related brain differences.

## Introduction

While sex-based brain differences are controversial, one of the most striking and well-accepted differences between males and females is their divergent susceptibility to certain neurological disorders^1,2^, for example, autism spectrum disorder (ASD)^3^ and schizophrenia^4–6^. This differential susceptibility points to fundamental differences in brain structure and/or function. However, the presence of clear, sex-specific neurological characteristics has remained a topic of heated debate, particularly since adult sex-based differences are not dimorphic, but rather a continuum with substantial overlap among individuals^7^. Nonetheless, certain quantitative differences in size are well established, with adult males having on average larger intracranial and total brain volumes and increased grey matter in several brain regions^7,8^. The neonatal presence of size differences, even after correcting for birth weight^9,10^, speaks to a developmental origin. Yet, how these size differences arise, and whether they relate to sex-biased susceptibility to neurological conditions, remain to be determined.

Sexual differentiation has two main sources: the cell-intrinsic chromosomal complement and circulating sex hormones^11^. The effects of sex hormones on brain development have been studied more extensively in mice^12^, where testosterone secreted by the embryonic testes was shown to have its masculinising effects through its conversion to estradiol (the predominant form of estrogens). Thus, counter intuitively, it is estradiol, and not testosterone, which masculinises the mouse brain^13^. However, the binary nature of rodent brain sexualisation and behaviour is strikingly different from humans, and thus, whether such a mechanism is at play in humans is unknown. Furthermore, studies of the cell-biological effects of sex hormones on embryonic neural progenitors^14,15^ have, so far, not been able to provide a mechanistic understanding of size differences, particularly in the human context.

## Results

### Androgens increase the number of basal progenitors in developing human neocortex

To gain an understanding of developmental events that may lead to male-female differences in the human context, we used cerebral organoids^16^, which provide a reliable model of early human brain development^17^. We compared male and female organoids, assessed neural progenitor responses to different sex hormones, and analysed the behaviour of cells and their progeny over time. Overall, we saw no differences between organoids made from female (XX) and male (XY) cell lines in various assays (see **Methods**), suggesting intrinsic sex chromosomal makeup does not influence cortical development, at least in this model system (Fig. 1, Extended Data Fig. 1–2).

**Figure 1.**
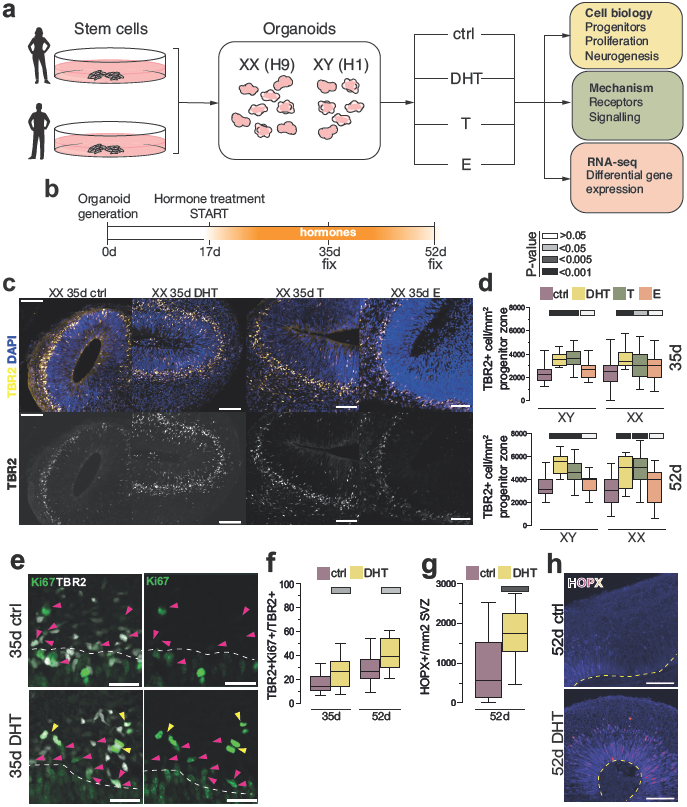
Androgens, but not estrogen, cause an expansion of progenitors. a) Schematic of the project workflow. b) Organoid generation and treatment protocol timeline. c) Female (XX) organoids at 35d, treated with DHT, T and E and stained for the intermediate progenitor marker TBR2 (yellow, white). Note the increase in TBR2+ in DHT/T treated organoids. d) Quantification of TBR2+ cells per mm^2^ of progenitor layer in male (XY) and female (XX) organoids, at 35d and 52d. e) Image of the ventricular zone/subventricular zone (VZ/SVZ) border in XX 35d organoids stained for Ki67 (green) and TBR2 (white). Cells positive for both Ki67 and TBR2 are indicated with magenta arrowheads. Double positive mitotic cells are indicated with yellow arrowheads. A white dashed line delineates VZ/SVZ border with the ventricular zone below. f) Quantification of Ki67/TBR2 double positive cells, out of all TBR2+ cells, in XX 35d and 52d organoids. g) Quantification of HOPX+ basal radial glia per mm^2^ SVZ of XX 52d organoids. h) Immunostaining for HOPX (fire LUT) of XX 52d organoids. Yellow dashed line indicates the ventricular surface. Note the HOPX signal in radial glia. All images are single, 1.2μm optical planes. Scale bars: c) and j) 50μm, g) 25μm. Whiskers in boxplots are min-max values. Significance values (Mann-Whitney, two tailed) of the treatment, as compared to the control, indicated by different shades of grey.

Next, to examine the effects of sex steroids, we exposed brain organoids to male sex hormones testosterone (T) and dihydrotestosterone (DHT), and the female hormone estradiol (E2) (Fig. 1a, b). DHT is a more potent androgen and cannot be aromatised into E2, as opposed to T^18^. Treatments were started at day 17 of the protocol as the organoids at this stage morphologically resemble the time point at which the testosterone surge begins in the human embryo (around 8 gestational weeks (GW))^19^.

In both male and female cell lines, the most striking phenotype was seen upon treatment with androgens (DHT or T), which led to an increase in intermediate progenitors, a population of basally located cells that undergo a finite number of divisions to produce neurons (Fig. 1c, d, Extended Data Fig. 1a-b). Notably, this phenotype was not seen upon treatment with E2. Staining for the proliferative marker Ki67 revealed an increased proportion of intermediate progenitors that remained proliferative upon DHT treatment (Fig. 1e, f). In addition, we observed an increase in the number of another basal progenitor type known as basal radial glia^20,21^ in DHT-treated organoids (Fig. 1g, h). Taken together, these findings suggest that androgens increase the numbers and proliferation of basal progenitors in the developing human neocortex.

### DHT promotes proliferation of apical radial glial stem cells

To investigate the mechanism of this increase in basal progenitors, we examined the neural stem cells that give rise to basal progenitors and neurons, called radial glia. We observed certain features indicative of radial glial proliferation, namely number of apical mitoses and ventricular length, that appeared increased upon treatment with androgens (Extended Data Fig. 2a, b). We next used viral sparse labelling of the radial glial neural stem cells to perform lineage tracing (Fig. 2a), and observed a significant increase in clone size upon treatment for 8 days with DHT (Fig. 2b, c, e). We then tested whether this effect was specific and reversible by removing hormone and labelling a novel population of radial glia with a different virus (Fig. 2a). After 8 days of no treatment, newly-labeled clone sizes of control and previously-DHT treated organoids were comparable (Fig. 2d, e). These results indicate that androgen signalling acts specifically and reversibly to promote symmetric, proliferative divisions of radial glial stem cells, increasing their output.

**Figure 2.**
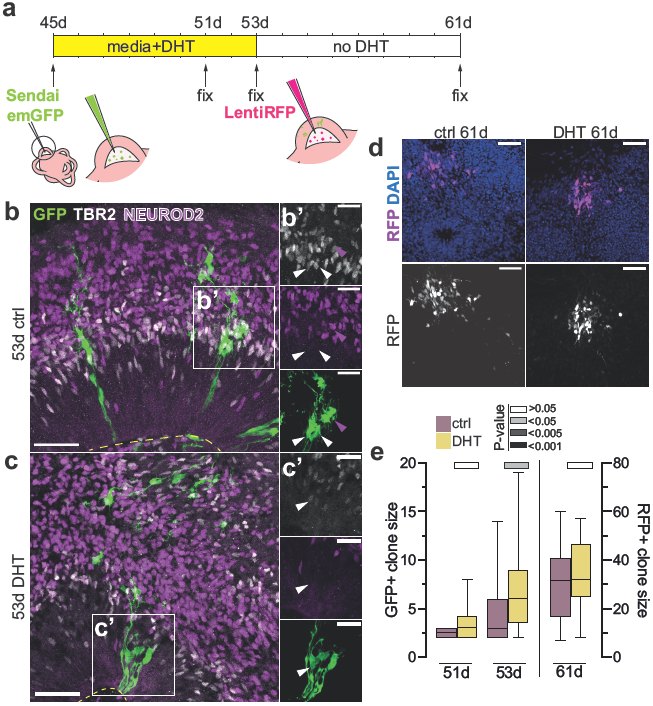
Androgens influence the mode of division of radial glia. a) Timeline and schematic of the Sendai/Lenti virus lineage tracing and analysis, with virus encoding emGFP injected at day 45 and virus encoding RFP injected at day 53. b) XX control organoids, injected with Sendai emGFP at 45d, fixed at 53d and stained for GFP (green), TBR2 (white) and NEUROD2 (magenta). Yellow dashed line indicates ventricular surface. b’) Inset from b). White arrowheads: TBR2/GFP double positive cells. Magenta arrowhead: NEUROD2/GFP double positive cell. Image is a maximum intensity projection of 9 individual 1.2μm-thick single optical sections. c) XX DHT-treated organoids, injected with Sendai emGFP at 45d, fixed at 53d and stained for GFP (green), TBR2 (white) and NEUROD2 (magenta). Yellow dashed line indicates ventricular surface. Image is a maximum intensity projection of 11 individual 1.2μm-thick single optical sections. c’) Inset from c). White arrowhead: TBR2/GFP double positive cell. Note the distribution of the clone, with most of the GFP+ cell bodies still in the VZ. d) XX organoids, injected with Sendai emGFP at 45d, with Lenti RFP at 53d and fixed after a pulse of DHT treatment or vehicle control. Immunostaining for RFP (magenta, white). Co-stained with DAPI (blue). Images are a maximum projection of 13 (control) and 11 (DHT) individual 1.2μm-thick single optical sections. e) Quantification of GFP+ clone size (left side) at 51d and 53d and RFP+ clone size at 61d. Individual data points represent individual clones. Scale bars: b), c), d) 50μm, b’), c’) 25μm. Whiskers in boxplots are min-max values. Significance values (Mann-Whitney, two tailed) of the treatment, as compared to the control, indicated by different shades of grey.

### Androgens signal through the androgen receptor expressed by radial glial stem cells

Androgen receptor (AR), the canonical androgen signalling mediator, is a nuclear receptor which, upon ligand binding, homodimerizes and translocates to the nucleus, where it acts as a transcription factor^22^. AR is present in the developing primate cortex^23^, but it is still unclear in which cells of the developing human neocortex. We performed RNAScope^24^ for AR mRNA, which revealed widespread signal in radial glia (Fig. 3a), with a distribution that varied depending on the stage of the cell cycle (Extended Data Fig. 3a, b). Western blot for AR protein revealed a decline over time, whereas organoids exposed to DHT showed a sustained expression of AR protein (Extended Data Fig. 3c). This is in accordance with similar observations in a neural progenitor cell line^15^. Together, these findings indicate that radial glial stem cells are receptive to androgens, and that androgens in turn positively feedback to stabilise AR protein^25^.

**Figure 3.**
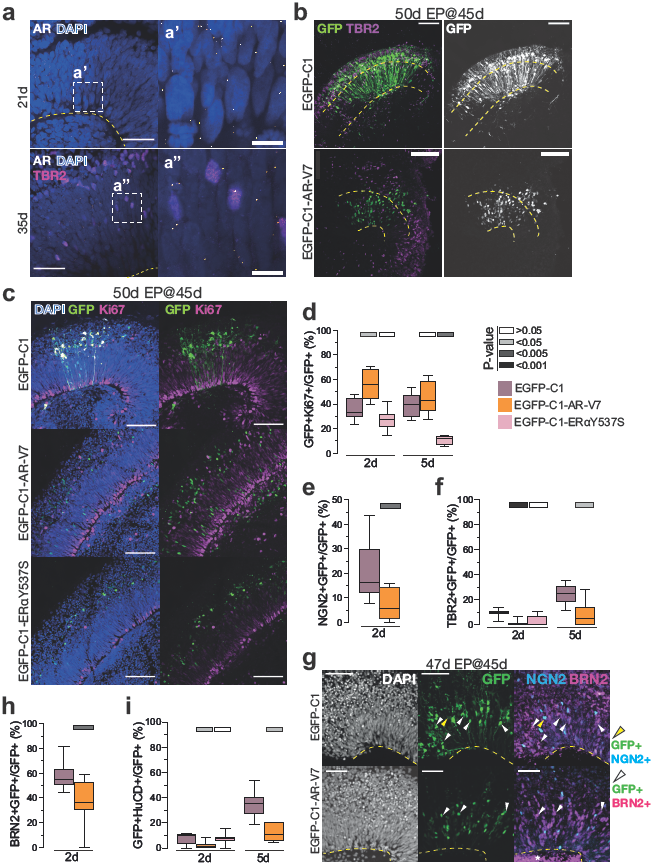
Androgen receptor signalling promotes proliferative identity of radial glia. a) RNA Scope for AR of XX 21d (top) and 35d (bottom) organoids. Single AR mRNA puncta (white). Immunostaining for TBR2 (magenta). Co-stained with DAPI (blue). a’) Portion of the ventricular zone (VZ) from XX 21d organoid. a”) Portion of the VZ from XX 35d organoid. b) Images of XX 35d organoids electroporated at 45d and fixed at 50d. EGFP-C1: control plasmid; EGFP-C1-AR-V7: plasmid expressing constitutively active AR. Immunostaining for GFP (green, white) and TBR2 (magenta). Yellow dashed lines demarcate the apical and basal boundaries of the VZ. Note the localization of GFP+ cell bodies in the VZ in EGFP-C1-AR-V7 electroporated organoids. c) Immunostaining for GFP (green) and Ki67 (magenta) on XX 50d organoids, electroporated at 45d. Co-stained with DAPI (blue). EGFP-C1: control plasmid; EGFP-C1-AR-V7: plasmid expressing constitutively active AR; EGFP-C1-ERαY537S: plasmid expressing constitutively active ERα. d) Quantification of the proportion of GFP+ cells costaining for the proliferation marker Ki67 2- and 5 days post electroporation in XX organoids electroporated at 45d. e) Quantification of the proportion of GFP+ cells costaining for the neurogenic marker NGN2 2 days post electroporation in XX organoids electroporated at 45d. f) Quantification of the proportion of GFP+ cells costaining for the intermediate progenitor marker TBR2 2- and 5 days post electroporation in XX organoids electroporated at 45d. At 5 days post electroporation, most cells electroporated with EGFP-C1-ERαY537S died. g) XX 47d organoids, electroporated at 45d. Immunostaining for GFP (green), NGN2 (cyan) and BRN2 (magenta). Co-stained with DAPI (white). Yellow arrowheads: GFP+/NGN2+ cells. White arrowheads: GFP+/BRN2+ cells. h) Quantification of the proportion of GFP+ cells costaining for the upper layer neurogenesis marker BRN2 2 days post electroporation in XX organoids electroporated at 45d. i) Quantification of the proportion of GFP+ cells costaining for the neuronal marker HuC/D 2- and 5 days post electroporation in XX organoids electroporated at 45d. At 5 days post electroporation, most cells electroporated with EGFP-C1-ERαY537S died. Scale bars: b), c), g) 50μm, a) 25μm, a’), a”) 5 μm. Whiskers in boxplots are min-max values. Significance values (Mann-Whitney, two tailed) of the treatment, as compared to the control, indicated by different shades of grey.

To test whether androgens act through AR within radial glia, we used a constitutively active form of AR, AR-V7^26^, which enters the nucleus and exerts its function, even in the absence of androgens. GFP tagged AR-V7 showed nuclear distribution, indicating its successful entry into the nucleus (Fig. 3b). Most of these GFP-positive nuclei resided in the apical ventricular zone at 2 days post-electroporation, indicating a radial glial identity, whereas in control organoids, cells expressing GFP alone were distributed in both apical and more differentiated basal compartments (Fig. 3b). AR-V7-expressing cells were healthy and cycling, as indicated by their Ki67 immunostaining (Fig. 3c), which also revealed a ~60% increase in proliferation compared with control (Fig. 3d). Along these lines, staining for markers of differentiation revealed a decrease in differentiation of cells expressing AR-V7 (Fig. 3b, e-g).

The AR signalling pathway is most heavily studied in the context of prostate cancer, where downstream signalling can lead to changes in the levels of the transcription factor BRN2^27^. BRN2 is present in primate radial glia and is important for neurogenesis^28^. Mirroring the processes described in prostate cancer, we observed a significant reduction in BRN2 in AR-V7-electroporated cells (Fig. 3g, h) further supporting a less differentiated state. Finally, AR-V7 expressing cells also exhibited a significant reduction of the neuronal marker HuC/D, indicating that they did not become neurons at the same rate as control cells (Fig. 3i).

In order to test the specificity of the effect of expression of AR-V7, we tested expression of a constitutively active form of another sex hormone receptor, estrogen receptor α (ERα)^29,30^. Expression of a GFP tagged constitutively active ERαY537S revealed proliferation status similar to the control at 2 days post-electroporation (Fig. 3c-d), indicating that the effects of AR-V7 were not due to a general response to the constitutive activation of any sex hormone receptor, but were AR specific. We did however observe that at 5 days postelectroporation most of the ERαY537S+ cells died (Fig. 3d), indicating a later effect on cell survival.

To directly visualise the effects of constitutively active AR, we performed live imaging of organoids electroporated with a control mCherry plasmid and plasmid encoding GFP-tagged AR-V7. As expected, control mCherry-only positive cells divided asymmetrically resulting in one daughter cell remaining a radial glial stem cell, while the other differentiated and left the apical ventricular zone (Extended Data Fig. 4). On the other hand, GFP tagged AR-V7 positive cells divided symmetrically with both daughter cells remaining in the ventricular zone, indicating a radial glial identity (**Extended Data Movie 1-3**). This behaviour was observed at two different organoid ages, even at 53 days, when most divisions of radial glial stem cells would otherwise be expected to be asymmetric, differentiative divisions.

Together, these findings reveal that when AR signalling is active, radial glial stem cells remain proliferative and increase their numbers. However, our earlier findings of an increased abundance of basal progenitors upon androgen treatment revealed that radial glia in the presence of androgens could still differentiate and produce basal progenitors. These findings at first glance appear contradictory; however, these two experimental approaches accomplished very different things. Whereas the AR-V7 isoform is constitutively active, and therefore would continuously signal in radial glia, application of androgens would only activate AR signalling in those cells that already express AR. Our findings of fluctuating cellular levels of AR (Extended Data Fig. 3b) suggest normally transient responsiveness of these cells, which, during an unresponsive phase, would have a chance to differentiate and generate basal progenitors. AR signalling would therefore be expected to increase the stem cell pool such that upon differentiation, they would produce increased numbers of basal progenitors.

### RNA-seq analysis of treated organoids reveals genes involved in androgen response

Androgen induced changes in gene expression in neural progenitors have previously been reported to be subtle^15^. Therefore, in order to capture the downstream response with enough transcriptomic coverage to detect such subtle changes we performed RNA-seq on total mRNA isolated from control and androgen-treated organoids (Extended Data Fig. 5a, b). To detect differentially expressed genes, we employed a recently described novel method^31^ designed to ensure high sensitivity while controlling the false discovery rate (see **Methods**).

Comparing control and androgen treated organoids we found 65 up-regulated and 51 down-regulated genes (Fig. 4a, Extended Data Table 1). Within the up-regulated genes were several genes with roles in neural development/progenitors: HDAC2^32^, HDAC3^33^, YBX1^34^, EML1^35,36^, QARS^37^, PLK^38^. Among the upregulated genes, it was encouraging to find 5α reductase (SRD5A1), the enzyme that converts T into DHT. For this gene, we validated the DHT-induced increase on the protein level by quantification of subcellular SRD5A1+ puncta (Fig. 4b).

**Figure 4.**
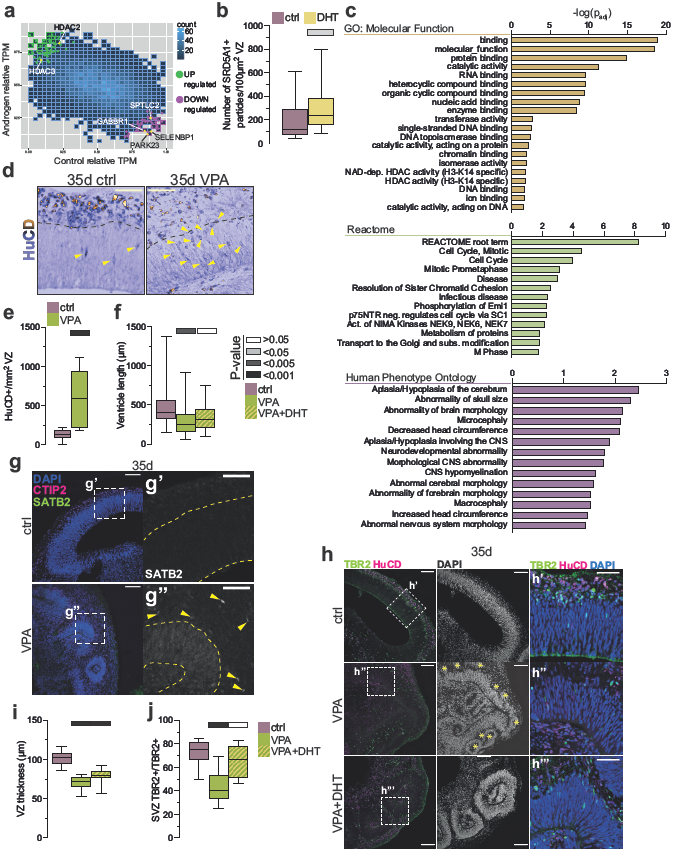
Differential gene expression shows involvement of HDACs in androgen-mediated phenotype. a) A plot showing normalised TPM values of all genes in control and androgen treatment. DEGs were detected using Delboy^31^. Upregulated DEGs are in green. Downregulated DEGs are in magenta. The density of data (count) is indicated by shades of blue. Among the downregulated genes, genes connected with schizophrenia are labelled. b) Quantification of SRD5A1, an upregulated DEG under androgen treatment. SRD5A1 was quantified as the number of SRD5A1+ puncta per 10μm^2^ of the ventricular zone (VZ) in XX 35d control and DHT-treated organoids (see Figure 5f). c) Graph of a number of top significant GO term enrichments in XX 35d androgen upregulated genes. d) HuC/D immunostaining (purple) in control and VPA-treated XX 35d organoids. e) Quantification of HuC/D+ cells per mm^2^ VZ in control and VPA-treated XX 35d organoids. f) Quantification of ventricular length in pm for XX 35d old organoids treated with VPA and VPA+DHT. g) Immunostaining for CTIP2 (magenta), and SATB2 (green, white) on XX 35d control and VPA-treated organoids. Co-stained with DAPI (blue). g’), g”) inset from g). g”) Note the SATB2+ cells (yellow arrowheads) and SATB2+ staining present in the VZ in VPA-treated organoids. Yellow dashed lines demarcate the apical and basal boundaries of the VZ. h) Immunostaining for TBR2 (green) and HuC/D (magenta) in XX 35d organoids, treated with VPA from 17d. Co-stained with DAPI (white, blue). h’) h”) h”’) inserts from h), as indicated. Yellow asterisks indicate neural rosettes forming above the VZ in some VPA-treated organoids. i) Quantification of the thickness of VZ in pm for XX 35d organoids treated with VPA and VPA+DHT. j) Quantification of the percentage of TBR2+ cells in the SVZ, out of all TBR2+ cells, for XX 35d organoids treated with VPA and VPA+DHT. Scale bars: h) 100μm, h’), h”), h”’), d), g) 50μm. Whiskers in boxplots are min-max values. Significance values (MannWhitney, two tailed) of the treatment, as compared to the control, indicated by different shades of grey.

Two of the upregulated genes, MCM4 and PHB, are listed in the Simons Foundation Autism Research Initiative (SFARI) gene list as strongly associated with ASD, (https://gene.sfari.org/, Category 3 or higher), and several others are involved in ASD^39–41^. 10 of the upregulated and 8 of the downregulated genes also appeared on the list of genes significantly differentially expressed in schizophrenia^42^. A hypergeometric test of the intersection between the datasets showed that both the overlaps (up- and downregulated genes) were significant (P=3.6E-4 and P=0.0012, respectively) (Extended Data Fig. 5c.).

Gene ontology analysis^43^ of downregulated genes showed very low enrichment scores, while upregulated genes were enriched in various terms associated with brain size, developmental disorders, and chromatin activity. In particular, GO: Molecular Function terms were connected to nucleic acid binding/chromatin binding and HDAC activity, Reactome terms were connected to cell cycle and M phase, and Human Phenotype Ontology terms were connected to neurodevelopmental disorders and CNS abnormality, particularly brain size and head circumference (Fig. 4c).

### Interfering with androgen signalling effectors influences differentiation

The effect on HDACs stood out in particular because dysregulated levels of HDACs have been suggested to be involved in masculinisation of the rodent brain^44^ and in pathogenesis of the male-biased disorders ASD^39,45,46^ and schizophrenia^47,48^, and several genes on the down-regulated list have previously been reported to be associated with schizophrenia (PARK23^49^, SELENBP1^50^, GABBR1^51^, SPTLC2^52^).

A potent inhibitor of HDACs, valproic acid^53^ (VPA), is commonly used as an antiepileptic, but prenatal exposure can lead to malformations of the embryo^54^. *In utero* exposure to VPA has been associated with increased risk of developing ASD^55^, potentially by its interference with the normal balance of proliferation and differentiation of neural progenitors^56^. We therefore tested for a functional interaction between HDACs and androgens by treating organoids with VPA. We added VPA at 0.7mM, which is close to therapeutic doses of VPA^57^. VPA-treated organoids had smaller ventricles and an increased number of neurons in the apical ventricular zone upon VPA treatment, as indicated by HuC/D staining (Fig. 4d-f), suggesting that HDAC inhibition promotes depletion of the radial glial stem cell pool and premature neuronal differentiation, and indicating that HDACs, like androgens, normally inhibit differentiation of radial glial stem cells. Furthermore, staining for later-born upper layer neurons revealed a premature production of these neurons upon VPA treatment (Fig. 4g), suggesting a premature switch in neurogenic potential, similar to previously shown HDAC2^58^ and HDAC3^33^ knock-out studies.

To test whether HDACs acted downstream of androgen signalling, we investigated whether organoids treated with VPA also exhibited phenotypes affecting progenitor populations. In addition to the smaller ventricular length, we observed a noticeably thinner ventricular zone (Fig. 4h, i) and a reduced number of intermediate progenitors (Fig. 4h, j). While administration of DHT together with VPA did not rescue the thickness of the ventricular zone in VPA-treated organoids (Fig. 4i), it did rescue the reduced ventricular length and returned intermediate progenitor distribution to control levels (Fig. 4h, j). This ability of DHT to rescue aspects of the VPA-induced phenotype, namely those processes also affected by androgen signalling, suggests that HDAC activity functions downstream of androgen signalling in cortical neurogenesis.

### Neural progenitors of the ventral forebrain respond to DHT differently

The finding that a number of the differentially expressed genes were associated with both ASD and schizophrenia points to a role for androgens in the sex-bias of these conditions^47–49^. There is increasing evidence in both ASD and schizophrenia for an imbalance in excitatory and inhibitory neurons^59,60^. Interestingly, excitatory and inhibitory neurons have distinct developmental origins. While excitatory neurons are produced by radial glial stem cells within the cortex, inhibitory neurons are generated in the ventral forebrain and later migrate into the cerebral cortex from afar^61,62^ (Fig. 5a). Because of these distant origins, we hypothesised that circulating androgens may impact inhibitory neural progenitors in the ventral telencephalon differently to precursors of excitatory neurons in the dorsal cortex. We therefore treated ventral forebrain organoids with sex steroid hormones and found that DHT actually caused a slight decrease in the number of ventral intermediate progenitors (Fig. 5b, c). This effect was androgen specific since E2 treatment had no effect. This indicates that different parts of the brain respond to androgens in markedly different ways, which could have significant implications for the final neuronal composition of the cortex.

**Figure 5.**
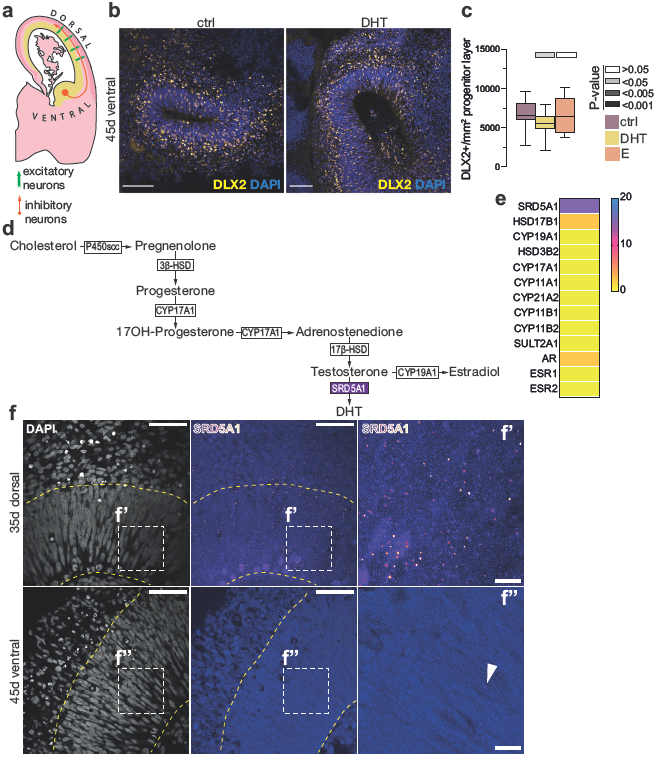
Androgens do not promote expansion of ventral progenitors. a) Schematic of neurogenesis in the human developing brain at ~12GW. Excitatory neurons (green arrows) are born within the dorsal cortex and migrate basally to form the cortical plate. Inhibitory neurons (orange arrow) are born in the ventral telencephalon and migrate towards the dorsal side to incorporate themselves into the cortical plate. Yellow: progenitor zone, dark pink: cortical plate. b) Immunostaining for a ventral intermediate progenitor marker DLX2 (yellow) on XX 45d ventral brain organoids. Co-stained with DAPI (blue). c) Quantification of DLX2+ cells per mm^2^ progenitor layer for control, DHT and E-treated organoids. d) Schematic of the steroidogenic pathway leading to the formation of DHT and estrogen (estradiol). SRD5A1 is indicated in purple. e) Heatmap of steroidogenic enzyme expression from XX 35d RNA-seq. Values are in tmp. f) Comparison of SRD5A1 (fire LUT) immunostaining on XX dorsal 35d and ventral 45d organoids. Co-stained with DAPI (white). Yellow dashed lines demarcate the apical and basal boundaries of the VZ. f’) Portion of the VZ in XX dorsal 35d organoids stained for SRD5A1. Note the SRD5A1+ puncta. f”) Portion of the VZ in XX ventral 45d organoids stained for SRD5A1. A white arrowhead indicates an SRD5A1+ punctum. Scale bars: f) 50μm, f’), f”) 10μm. Whiskers in boxplots are min-max values. Significance values (Mann-Whitney, two tailed) of the treatment, as compared to the control, indicated by different shades of grey.

Importantly, the finding that DHT is capable of eliciting these androgen responses in neural progenitors suggests that, because DHT cannot be aromatized, these effects must not be through its conversion to estradiol, unlike in rodents^13^. In support of this conclusion, we found that aromatase (CYP19A1), the enzyme that converts testosterone into estrogen^18^ (Fig. 5d), was absent in our RNA-seq dataset (Fig. 5e), whereas SRD5A1^18^, the enzyme that locally converts T to DHT^63^, was abundantly expressed. Furthermore, staining for SRD5A1 revealed punctate signal in neocortical regions of dorsal organoids, primarily in radial glia, but almost no SRD5A1 puncta in ventral organoids (Fig. 5f). This difference in SRD5A1 presence indicates that these two brain regions (dorsal and ventral) might have a different natural aptitude to respond to androgens during *in vivo* brain development.

### Additional progenitors lead to an increased number of neurons

The enrichment for ontology terms associated with brain size (Fig. 4c), as well as the fact that males on average exhibit a larger brain size, pointed to a potential role for effects of androgens on brain size and neuron number. However, we did not observe any change in neuronal numbers upon androgen treatment despite the increase in basal progenitors (Fig. 6a-c). We hypothesised that, due to the continuous nature of these treatments, the progenitors were maintained in a proliferative state. We therefore performed a pulse-chase experiment in which organoids were subjected to hormone for 18 days (Fig. 6d), followed by their removal from the media until fixation 17 days later. Analysis of neuronal composition showed a subtle, but significant increase in NEUROD2+ neurons upon DHT treatment (Fig. 6e). However, staining for the subset of neurons that make up the early-born deep-layers using the marker CTIP2 revealed no change (Fig. 6f, g). We hypothesised that this may be due to androgen-mediated maintenance of progenitor proliferation during the stage at which these neurons would normally be produced, whereas later born neurons may be expected to show an increase upon their “release” from this proliferative state, after the removal of androgen. Indeed, staining for the upper-layer marker, SATB2, revealed a striking increase in these later born neurons (Fig. 6h, i). These findings demonstrate that an androgen-induced switch towards more proliferative divisions of neural progenitors results in a later increase in excitatory neuron numbers within the neocortex.

**Figure 6.**
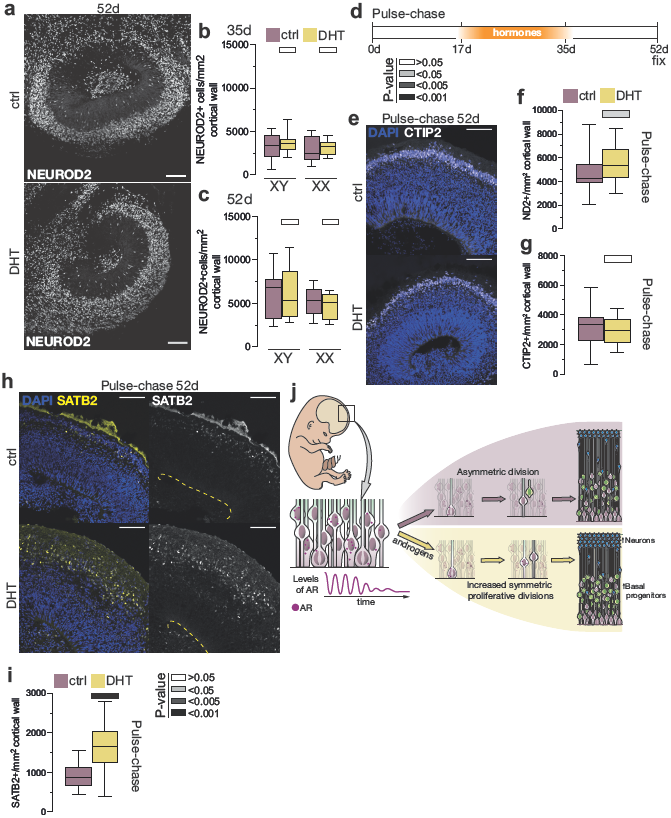
Androgens cause an increase in neuronal numbers. a) Immunostaining of XX 52d, control and DHT-treated, organoids stained for NEUROD2 (white). b) Quantification of NEUROD2+ cells per mm^2^ of cortical wall at 35 days, in XY and XX organoids. c) Quantification of NEUROD2+ cells per mm^2^ of cortical wall at 52 days, in XY and XX organoids. d) Timeline of the pulse-chase experiment. Hormones were administered between 17-35d, then washed out and cultured in media alone. Organoids were fixed at 52d. e) Quantification of NEUROD2+ cells per mm^2^ cortical wall after pulse-chase treatment. f) Immunostaining for CTIP2 (white) of XX 52d organoids after pulse-chase treatment (control and DHT). Co-stained with DAPI (blue). g) Quantification of CTIP2+ cells per mm^2^ cortical wall after pulse-chase treatment. h) Immunostaining for SATB2 (yellow, white) of XX 52d organoids after pulse-chase treatment. Co-stained with DAPI (blue). Yellow dashed line indicates the ventricular surface. i) Quantification of SATB2+ cells per mm^2^ cortical wall after pulse-chase treatment. j) A proposed model of the effects of androgens on the developing neocortex. Androgen receptor is present in radial glia during embryonic brain development, but oscillates during the cell cycle and its levels progressively get lower as development progresses. At the onset of neurogenesis, androgens are secreted by the male embryo’s testes and act on the division mode of radial glia. Radial glia under the influence of androgens exhibit increased propensity to undergo symmetric proliferative divisions, rather than asymmetric ones, hence increasing the stem cell pool. With the levels of AR dropping over time, effects of androgens weaken and an increased pool of radial glia gives rise to an increase in basal progenitors. Finally, an overall increase in basal progenitors results in more excitatory neurons. Scale bars: a), h) 100μm, e) 50μm. Whiskers in boxplots are min-max values. Significance values (Mann-Whitney, two tailed) of the treatment, as compared to the control, indicated by different shades of grey.

## Discussion

Because sex-based brain differences have remained controversial, we sought to provide some clarity to this issue by examining this question from a cell biological and mechanistic standpoint. We examined whether sex-based differences could be observed during human brain development using cerebral organoids as a model system. Although even the most well-established sex-based brain differences are subtle, we were able to observe significant differences in neural progenitor numbers and division modes under the influence of androgens, that overall suggest a shift towards proliferative divisions of radial glial stem cells. These increased stem cells are then able to produce a higher number of basal progenitors, and eventually increased neuron numbers (Fig. 6j).

The fact that we saw highly similar effects of androgens in both male and female brain organoids points to primarily cell-extrinsic driven sex differences in cellular response at this stage of development. This would also fit with the non-binary nature of brain sex-related phenotypes, since a more quantitative (rather than qualitative) difference would emerge from changes in concentrations and timing of sex hormones. Such fine-tuning may also come from the responsiveness of the cells, and our findings of transitory expression of AR in radial glia would support that notion. Thus, while androgens seem to have a general proliferative effect on neural progenitors of the dorsal neocortex, the subtle, cell stage-dependent, and dynamic nature of the androgen response calls for future studies.

In rodents, androgens themselves are not thought to impact neurogenesis during development, although during adult neurogenesis androgens were shown to promote survival of new neurons^64^. Instead, it is estradiol that influences rodent neural progenitors^14^ and masculinises the brain^13,65,66^. However, we observed that in human brain organoids, administration of estradiol caused only minor changes. This discrepancy indicates that either estradiol causes phenotypes that we were not able to observe using this set of parameters, or the timing of these studies is not appropriate for investigation of the role(s) of estrogens. It could also point to human, or perhaps more widely primate, specific biological processes that require further investigation.

The identified androgen regulated genes point to chromatin remodelling and HDAC activity, and our functional studies further support the role of HDACs in mediating the effects of androgens on neural progenitors of the developing human neocortex. HDAC activity has been linked to ASD and schizophrenia, both of which also exhibit a sex-bias and present with sex-differential manifestations. While prenatal sex hormone levels have been primarily examined in connection with ASD^15,67^, some reports have shown an association of prenatal androgens and the development of schizophrenia^68^. Our findings connect these risk factors, and suggest that the activity of androgens on HDACs in the developing human neocortex, through their previously established pro-proliferative effects^33,58,69,70^, may help explain the sex-bias of these disorders.

Sex biased neurological and mental health disorder susceptibility suggests the presence of morphological and/or functional differences between healthy male and female brains that may predispose to certain conditions. The most conclusive differences are quantitative in nature, namely brain size ^7,9,10,71^, and our findings of increased cortical excitatory neurons upon androgen treatment would suggest that such differences may, at least in part, arise due to developmental regulation of progenitor behaviour. Interestingly, we did not observe such an effect of androgens in the area that produces inhibitory neurons, pointing to an important role in regulating the balance of excitatory to inhibitory neurons. Along these lines, ASD and schizophrenia seem to share a pathophysiological mechanism of excitatory to inhibitory neuron imbalance ^59,72–74^. Thus, our findings are able to consolidate these various observations and suggest that male sex hormones act through HDAC activity to regulate excitatory neuron number and hence brain size. Misregulation of this process could affect the balance between the numbers of excitatory and inhibitory neurons and thus may impact risk for developing sex-biased disorders such as ASD and schizophrenia.

## Materials and Methods

### Stem cell maintenance and organoid generation

Male (XY) and female (XX) stem cells (male - H1 (WiCell WA01) and female - H9 (WiCell WA09)) were approved for use in the project by the UK Stem Cell Bank Steering Committee. Cells were cultured on Matrigel^®^ (Corning, 356234) – coated plates and maintained in StemFlex™ medium (Thermo Fisher, A3349401). We used the commercial kit (STEMdiff™ Cerebral Organoid Kit, StemCell Technologies 08570) with previously described modifications to increase embryoid body surface area^75^, namely microscaffolds (18000 cells seeded) or smaller EBs (2000 cells seeded). Organoids were transferred to the orbital shaker at 57rpm (ɸ25cm) at 15d (±1 day, depending on the morphology). Matrigel^®^ was removed manually between days 14-17, after visual inspection. Organoids were transferred to IDM+A supplemented with diluted Matrigel^®^ on day 30 and kept at those conditions until fixation. Ventral organoids were generated as previously described^76^, with ventralizing factors (2.5μM IWP-2, Sigma, I0536; 100nM SAG, Sigma, 566661) supplemented in the neural induction (NI) medium for 2 days.

### Hormone and drug treatments

All steroid hormones were diluted in 96% ethanol (EtOH) to obtain stock solution. Organoids were treated with 30nM or 100nM dihydrotestosterone (DHT, Sigma A8380-1G), 100nM testosterone (Sigma, T1500), 100nM estradiol (active form of estrogen) (Sigma, E1024) and 100nM progesterone (Sigma, P8783). 96% ethanol was used as a vehicle control (1.4μl of 1:10 ethanol in PBS to 5mL of media, which does not impact the normal development of the neuroepithelium). The treatments started on day 17 of the protocol. Hormone treatments were administered twice a week, with changing of the media. Organoids were collected for analysis at 35 and 52 days. After establishing the phenotype in both cell lines, and since we observed no significant difference between cell lines in their response to different hormones (two-way ANOVA *P*=0.23), we focused on the female cell line for subsequent experiments, as it showed greater batch-to-batch consistency.

Organoids exposed to 100nM DHT for prolonged periods of time exhibited impaired morphology at later stages (increased cell death and general cytoarchitectural disturbances, data not shown), so we reduced the amount of DHT to 30nM. This amount was chosen because the potency of DHT is roughly three times that of testosterone^77^, meaning any effects caused by 100nM T and 30nM DHT would be comparable.

The application of progesterone to the organoids did not yield any observable phenotype (data not shown). This also eliminated the possibility that progesterone present in the commercially available media (present at 20nM in N2 and B27 supplements, which are routinely added to the organoid media) might have an effect on organoid development.

For continuous hormone treatments (Sendai/Lenti virus labelling experiments), hormone treatments started immediately after labelling (day 30 or day 45) and were administered every day with the change of media.

Treatment with valproic acid (VPA) (sodium valproate; Stratech, S1168-SEL-200mg) at 0.7mM was started at 17d and continued through until 35d, with fresh drug supplementation twice a week during media change. DHT was added at 30nM in the VPA+DHT experiment. Organoids were collected for analysis at 35d.

The pulse-chase experiment was started at 17 days, and hormones were removed from the media at 35 days. Organoids were then grown in IDM+A with dissolved Matrigel^®^ and fixed at 52 days.

### Sendai and Lenti virus labelling

Organoids were labelled using a CytoTune™ EmGFP Sendai Fluorescence Reporter (ThermoFisher, A16519). The stock solution of the virus was diluted with PBS to 1:10 and kept on ice. The solution was visualized with Fast Green FCF (Sigma, F7252) and was delivered to the ventricles at 45d using a pulled capillary. Organoids were immediately subjected to hormone treatment. The media was changed every day. Organoids were collected for analysis at 6- and 8-days post-labelling. At 8 days post labelling, some of the organoids were injected with LentiBrite™ RFP Control Lentiviral Biosensor (Millipore, 17-10409), using the same procedure as with EmGFP Sendai virus. These organoids were kept in IDM+A with dissolved Matrigel^®^ for a further 8 days and then fixed for clonal analysis.

### Plasmids

Constitutively active androgen receptor (EGFP-C1-AR-V7)^26^ was obtained from Michael Mancini and Marco Marcelli (Addgene #86856). A plasmid containing estrogen receptor alpha (pEGFP-C1-ERα) was obtained from Michael Mancini (Addgene #28230) and mutated into a constitutively active form (pEGFP-C1-ERαY537S)^29^ using the Q5^®^ Site-Directed Mutagenesis Kit (New England Biolabs, E054S).

### Electroporations

Organoids were injected and electroporated at 45d as described previously^16^ with the following parameters: voltage − 40mV, duration – 50ms, pulses – 5, interval – 1.0s. All plasmids were used at 1μg/mL. EGFP-C1 plasmid was used as a control. Organoids were collected 2- and 5-days post-electroporation and media was changed at 2 days post-electroporation.

### Organoid fixation and processing

Organoids were collected at different stages and fixed in 4% PFA in phosphate buffer (PB) for 1 hour at room temperature (RT). After washing in PBS (2×), organoids were infiltrated with 30% sucrose in PB at 4°C until the tissue sank. After embedding in 7.5% gelatin/30% sucrose in PB and freezing in isopentane at −40°C, tissue blocks were stored at −80°C until sectioning. Organoids were sectioned using a cryostat (Leica, CM1950) at 20μm and sections were collected on coated slides (ThermoFisher Superfrost^®^ Plus, J1800AMNZ). Sections were stored at −20°C until further processing.

### Immunohistochemistry and RNA Scope

Immunostaining was performed as follows: Slides were briefly washed in PBS. Antigen retrieval (if necessary) was performed by using 0.01M sodium citrate, pH6.0. Slides were submerged in the antigen retrieval solution and heated for one hour at 70°C in a water bath. After antigen retrieval, slides were left to cool for 20 minutes at RT. After washing with PBS, sections were blocked with 0.25% Triton-X 100 with 4% donkey serum in PBS (permeabilization buffer) for 1-2 hours at RT. Primary antibodies were diluted in 0.1% Triton-X 100 with 4% donkey serum in PBS (blocking buffer) and left overnight at 4°C. The following primary antibodies were used: rat anti-histone H3 (phospho S28) (Abcam, ab10543, 1:500), rat anti-CTIP2 (Abcam, ab18465, 1:200), mouse anti-HuC/D (LifeTech, A21271, 1:200), mouse anti-Ki67 (Dako, MIB-1, 1:100), mouse anti-DLX2 (Santa Cruz, sc-393879, 1:200), mouse anti-SATB2 (Abcam, ab51502, 1:50), rabbit anti-NEUROD2 (Abcam, ab104430, 1:200), rabbit anti-Ki67 (Abcam, ab15580, 1:200), rabbit anti-NGN2 (Cell Signalling Technologies, D2R3D, 1:200), rabbit anti-HOPX (FL-73) (Santa Cruz, sc-30216, 1:100), chicken anti-GFP (Thermo Fisher, A10262, 1:500), goat anti-SRD5A1 (Abcam, ab110123, 1:200), goat anti-BRN2 (Santa Cruz, sc-6029, 1:200), sheep anti-TBR2 (R&D Systems, AF6166, 1:200).

The next day, sections were washed with PBS and secondary antibodies coupled to Alexa Fluor fluorescent dyes (ThermoFisher Scientific) were added (diluted in blocking buffer 1:1000) in combination with DAPI (1:1000). Sections were incubated for 1 hour at RT. After PBS washes, slides were mounted in ProLong™ Diamond (ThermoFisher, P36961), left to dry at RT in the dark, and stored at 4°C after imaging.

### Analysis of AR mRNA

AR mRNA was detected using the RNAscope^®^ Fluorescent Multiplex Reagent Kit (ACD, 320850). The probe used was Hs-AR-O2 (ACD, 522941). The slides were dehydrated using a sequence of increasing concentrations of EtOH (50-100%). Dehydrated sections were air-dried for 5’ at RT. Heat mediated target retrieval was performed in Target retrieval solution for 2’ at 95°C. Diluted protease III (1:15) was applied and incubated for 15’ at 40°C in a humidified chamber. After washing 3 times in PBS, pre-heated probe was applied to the slides and hybridised for 2 hours at 40°C in a humidified chamber. After hybridisation, the slides were washed in wash buffer twice. AMP 1-FL was applied and incubated for 30’ at 40°C. Sections were washed twice for 2’ in 1x Wash buffer at RT. AMP 2-FL, AMP 3-FL and AMP 4-FL were individually added to the slides and incubated for 15’, 30’ and 30’ at 40°C, respectively, with wash steps in 1x Wash buffer in between (twice for 2’ at RT). The slides were washed 3 times in MilliQ water. Permeabilisation buffer (0.4% Triton-X, 4% normal donkey serum in PBS) was applied and incubated for 1 hour at RT. Primary antibody (diluted in blocking buffer) was incubated over night at RT in a humidified chamber. The following day, slides were washed 3 times for 5’ in PBS. Secondary antibody and DAPI were incubated for 2 hours at RT. Slides were washed in PBS 3 times for 5’ and mounted in ProLong™ Diamond.

For analysis of AR mRNA after cell division, XX organoids were injected with EmGFP Sendai Fluorescence Reporter at 50d as described above to trace individual radial glia stem cells and their progeny. Organoids were fixed at 54d and submitted to the RNA Scope protocol. The AR mRNA signal was analysed only in well-defined GFP+ pairs of cells. Only AR mRNA puncta that were inside the GFP signal were taken into consideration.

### Imaging and image analysis

Imaging was performed either by using a Panoramic Confocal slide scanner, or Zeiss 710, 780UV or 880 confocal microscopes. Objectives used: 10×, 20×, 40× and 64×. Images were taken as either Z-stacks, or individual planes. Quantification of data from all systems were compared to ensure reproducibility of the data. Images obtained were processed using Fiji^78^.

### Quantifications

All quantifications were done using Fiji. Manual cell counting was performed by a custom plug-in, written by Johannes Schindelin. Only ventricles of appropriate morphology (distinct progenitor and neuronal layers, see below) and healthy appearance were chosen for quantifications.

### Cell type, morphology and proliferation quantifications

The ventricular length was defined as an uninterrupted stretch of apical surface populated by apical progenitors, as shown by phosphohistone H3 (PH3) or an apical progenitor marker staining (PAX6 or SOX2, not shown). The ventricular zone (VZ) was defined as a layer of cells directly abutting the ventricle, with a distinctive radial organisation of nuclei and high density of cells, as revealed by DAPI staining. The subventricular zone (SVZ) was defined as a zone basally adjacent to the VZ, with cells expressing TBR2 and nuclei of a rounder shape. Progenitor zone was defined as the combined area of the ventricular (VZ) and subventricular zone (SVZ) expressing the progenitor marker TBR2. The cortical wall was defined as the area between the apical and the basal side of the cortical lobule. The cortical plate was recognised as a zone at the basal side of the cortical wall containing neurons arranged in an ordered fashion. All counts were normalized per mm^2^, unless indicated otherwise. Counts were done without using pseudocolour. Quantifications are represented as boxplots. The body of the boxplot represents the 25^th^ to the 75^th^ percentile of the data. The line represents the median. Whiskers represent the minimum and maximum. All quantifications were done on a minimum of three independent organoid batches, with a minimum of 5 ventricles per treatment, per batch. Only live cells were counted. When quantifying multiple stainings, the cell was scored as double positive only if the signal from both channels analysed clearly corresponded in shape.

### Clonal analysis

Only cells labelled GFP-(labelled with CytoTune™ EmGFP Sendai Fluorescence Reporter) or RFP-positive (labelled with LentiBrite™ RFP Control Lentiviral Biosensor) were analysed. During imaging, special care was taken to collect all the GFP- or RFP-expressing cells from the 20μm section in the form of a Z-stack. Only clones consisting of >2 cells were taken into account, to ensure that the cell had divided upon receiving the virus. Only isolated clones were counted, in order to minimize the probability of several founder cells being infected upon the delivery of the virus, and their clonal progeny intermingling. Measurements were done on a minimum of 3 independent organoid batches, with a minimum of 2 organoids per timepoint, with a minimum of 5 clones analysed per timepoint, per treatment, per batch.

### Quantification of SRD5A1 puncta

After immunostaining with anti-SRD5A1 antibody, images were obtained using a 63× objective, with exactly the same laser power, exposure time and digital gain for both the control and DHT-treated sections. A square measuring 100×100 μm was designated in the VZ. Threshold was adjusted in order to remove any immunostaining background which was easily discernible from the SRD5A1 signal. SRD5A1 puncta were then automatically counted using the “Analyze particles” option.

### Live imaging

Organoids were electroporated at day 31 or 54, as described above. Plasmids containing pCAG-mCherry and EGFP-C1-AR-V7 were mixed to 300ng/mL and 1μg/mL, respectively, and delivered to the ventricles. Electroporated organoids were transferred to IDM+A+Matrigel^®^ medium for several hours for recovery. After that, organoids were embedded in 3% Low melting agarose (Sigma, A9414) in HBSS (1×) (Gibco, 14025092), left to settle and sectioned on the vibratome at 300μm. Sections were placed on Millicell Organotypic Cell Culture Inserts (Merck, PICM0RG50) and supplemented with 800μl serum-free media^79^. After 24 hours at 37°C, slices were imaged using Zeiss 880 Airy Scan, with the following parametres: 20× objective, FAST scanning mode, 1072×1072 pixels, scanning interval of 10 - 11.45 minutes, total imaging time >25 hours. Movies were processed using Zeiss ZEN software and Fiji.

### Western blot

Organoids for Western blot were treated with 30nM DHT or 100nM E every day, starting from d17 and collected every two days. Organoids (7 to 10 per condition) were washed twice in PBS prior to snap-freezing in liquid nitrogen. XX stem cells were used as a positive control, and XX d75 organoids as a negative (due to very low AR expression at this stage). Organoids and cells were lysed in modified RIPA buffer (mRIPA: 1% Triton-X, 0.1% SDS, 150mM NaCl, 50mM Tris pH7.4, 2mM EDTA, 12mM sodium deoxycholate) supplemented with protease (Thermo Fisher, 78430) and phosphatase (Sigma-Aldrich, 4906845001) inhibitors. Quick Start Bradford Dye Reagent (Bio-Rad, 5000205) assay was used to determine protein concentration. 20μg of total protein was loaded for every sample except for the negative control where 10μg of protein were loaded. Samples were run on SDS polyacrylamide gel electrophoresis (SDS-PAGE) 4-20% Tris glycine gel at 90V in MOPS-SDS buffer and transferred overnight at 30V to Amersham Hybond P 0.45 PVDF blotting membranes (GE Healthcare, 10600023). Membranes were blocked in 5% BSA for 2 hours at RT and incubated overnight with the following primary antibodies, rabbit anti-AR (Cell Signalling, D6F11, 1:2000), mouse anti-GAPDH (Abcam, ab8245, 1:2000). Membranes were washed in TBST buffer prior to incubation with secondary fluorophore conjugated antibodies 1:2000 for 1h at RT (goat anti-mouse DyLight 680 Invitrogen A21058, goat anti-rabbit DyLight 800 CST 5151). Membranes were washed in TBST and imaged using a Li-COR Odyssey CLx Infrared Imaging System.

### RNA-Seq library preparation and sequencing

Organoids were generated from the XX cell line and treated as described above (hormone treatment started at 17d). Organoids were collected at 35d, snap-frozen in liquid nitrogen and stored at −80°C until processing. Two to three individual organoids were collected per treatment. Total RNA from individual organoids was isolated using the RNeasy Mini Kit (Qiagen, 74104). Genomic DNA was removed using the Invitrogen™ TURBO DNA-*free*™ kit (Invitrogen, AM1907). After isolation, total RNA from individual organoids of the same treatment was pooled for each library preparation. RNA quality was assessed using Bioanalyzer (Agilent, G2939BA). Libraries were prepared using the NEBNext^®^ Ultra II DNA Library Prep Kit for Illumina^®^ (New England BioLabs, E7645) with starting input of 500ng of total RNA per sample. Ribosomal RNA was removed and mRNA isolated using the NEBNext^®^ Poly(A) mRNA Magnetic Isolation Module (New England BioLabs, E7490). During library preparation, AMPure XP Beads (Beckman Coulter, A63881) were used for reaction clean ups. NEBNext^®^ Multiplex Oligos (dual index primers) (New England BioLabs, E7600) were used in the PCR enrichment step, with 11 cycles. Library quality was assessed by Bioanalyzer and the concentration was determined using the NEBNextv Library Quant Kit for Illumina^®^ (New England BioLabs, E7630). Libraries were pooled and sequenced on the HiSeq4000 machine (Illumina) at 50 base pair length, single ended, to a minimum of 30 million reads per library.

### Data analysis and statistics

Analysis of cell number quantification was performed using GraphPad Prism 8 software. Statistical analyses of control *versus* hormone treatments were done as multiple pairwise analyses, using unpaired non-parametric test (Mann-Whitney, two-tailed) with significance threshold of P<0.05.

FASTQ sequence read files were aligned to the hg38 transcriptome (with decoy sequences) and quantified using salmon v0.14.1^80^ applying both a sequence bias correction and a GC bias correction with a mean fragment length of 300 bp and a standard deviation of 50 bp. Salmon quantification files containing Transcripts Per Million (TPM) estimates were imported into R using tximport v1.16.1 and DESeq2 v1.28.1^81^. Prior to analysis, a non-parametric empirical Bayesian batch-correction was applied to the data using the function ‘ComBat’ in sva v3.36.0^82^ (Extended Data Fig. 5a, b), and only genes with >4 TPM on average per sample (6 androgen (DHT/T), 3 control) were retained for downstream analysis.

Differential expression analysis was conducted using a novel approach^31^ that employs a logistic elastic-net regression implemented in the R package glmnet v4.0-2^83^ using an alpha of 0.5. The response variable was taken to be the sample hormone status (Androgen or Control), and the independent variable was the transcript abundance estimates for individual genes. To determine the value of lambda – the parameter controlling the strength of the regression penalty - for selecting differentially expressed genes, cross-validation was performed using the function ‘cv.glmnet’ and the ‘type.measure’ argument set to ‘deviance’. The value of lambda that minimised the deviance (a measure of goodness-of-fit) of the model was selected, and genes with a non-zero regression coefficient at the selected lambda value were considered to be differentially expressed, which included both up- and down-regulated genes. Log-fold change estimates for the selected genes were estimated using DESeq2.

Validation of the logistic elastic-net model was carried out using the same input dataset in which a ComBat batch-correction was applied to the treatment variable (Androgen or Control) to remove signal from the dataset. Then, a known amount of log-fold change signal was added to a random sample of genes using the binomial-thinning approach implemented in the R package seqgendiff v1.2.2^84^ using the function ‘thin_diff’ and randomly selecting 6 of the samples to be androgen samples and 3 to be controls. To determine the number of genes to add signal to, the R package locfdr v1.1-8^85^ was used to fit a mixture model to DESeq2 raw p-values derived from the original unmodified dataset. This package enables an estimate of the total number of non-null genes in the dataset – in this case 362. ‘locfdr’ was also used to infer the distribution of log-fold changes for non-null genes using DESeq2 log-fold change estimates derived from the original unmodified dataset. 362 log-fold changes were then randomly sampled from this distribution and paired with 362 randomly sampled genes. Both DESeq2 and the elastic-net models were then used to infer differential expression in this modified dataset and both the sensitivity and precision of each method was calculated. While DESeq2 achieved 100% precision, it had a sensitivity of just 2% (genes with an adjusted p-value < 0.1 were considered differentially expressed). In contrast, the elastic-net method achieved a sensitivity of 20% (72 correctly called genes) with a precision of 96% - all three false positives (out of 75 hits) had absolute log fold changes below 0.31.whereas 75% of the 72 true positive hits had an absolute log fold change above 0.37. In addition, false negatives had a significantly lower log fold change than true positives (Wilcoxon rank sum test, p = 3e-13). A reproducible R markdown script and data can be accessed at https://github.com/alextkalinka/hormone-DE-analysis.

## Acknowledgements

We would like to thank Deepak Srivastava, Sean Munro and Manu Hegde for helpful comments. We would like to thank the MRC LMB Light Microscopy facility and Bioinformatics (Paula Freire-Pritchett), as well as the CRUK Genomics facility. We would like to thank Michael Mancini, Marco Marcelli and Elizabeth Wilson for depositing their plasmids in Addgene. We thank the other members of the Lancaster lab and the MRC LMB Cell biology division for fruitful discussions. Work in the Lancaster lab is supported by the Medical Research Council (MC_UP_1201/9) and the European Research Council (ERC STG 757710).

## Author contributions

IK conceived the study, performed experiments, analysed data and wrote the paper. IC performed experiments and analysed the data. LP performed experiments. ATK analysed the RNA-seq data. MAL supervised the study and co-wrote the paper.

---

**Extended Data Figure 1.**
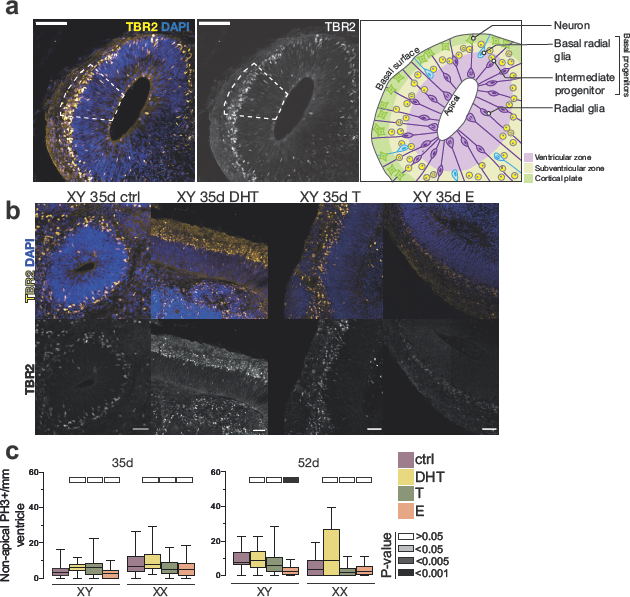
a) Illustration of the determination of progenitor zone for quantifications. The progenitor zone (VZ+SVZ) was determined based on immunostaining and histological landmarks (see Methods). Immunostaining of XX 35d organoids stained for TBR2 (yellow, white), co-stained with DAPI (blue). Dashed line indicates the progenitor zone. Schematic of an organoid ventricle with different progenitor populations of interest. b) XY 35d organoids stained for TBR2 (yellow, white), co-stained with DAPI (blue), treated with DHT, T or E. c) Quantification of non-apical mitotic, PH3+ cells in XY and XX, 35d and 52d organoids, treated with DHT, T and E. Scale bars: a), b) 100 μm. Whiskers in boxplots are min-max values. Significance values (Mann-Whitney, two tailed) of the treatment, as compared to the control, indicated by different shades of grey.

**Extended Data Figure 2.**
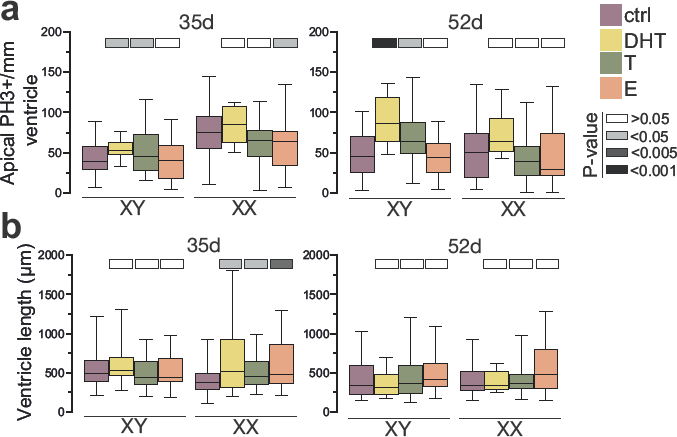
a) Quantification of apical mitotic cells (PH3+), normalized per mm length of ventricle in male (XY) and female (XX) organoids, at 35d and 52d, treated with DHT, T and E. b) Quantification of ventricular length in male (XY) and female (XX) organoids, at 35d and 52d, treated with DHT, T and E. Whiskers in boxplots are min-max values. Significance values (Mann-Whitney, two tailed) of the treatment, as compared to the control, indicated by different shades of grey.

**Extended Data Figure 3.**
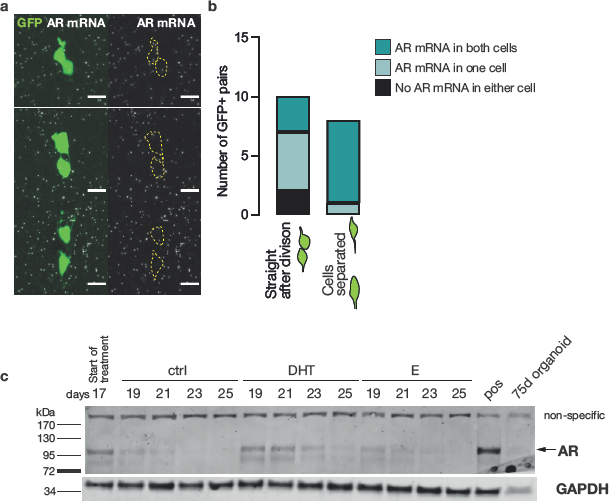
a) RNA Scope of androgen receptor (AR) mRNA (white puncta), together with GFP signal (green) from EmGFP Sendai Fluorescence Reporter showing examples of three daughter cell pairs. Yellow dashed lines indicate GFP+ cell body. b) Quantification of AR mRNA distribution in GFP+ pairs of cells, depending on the stage of cell division. c) Western blot for AR on control, DHT and E treated organoids at 17-25 days. AR specific band is as predicted to be ~110kDa. Note the increased AR signal in DHT-treated organoids. Scale bar: a) 5μm.

**Extended Data Figure 4.**
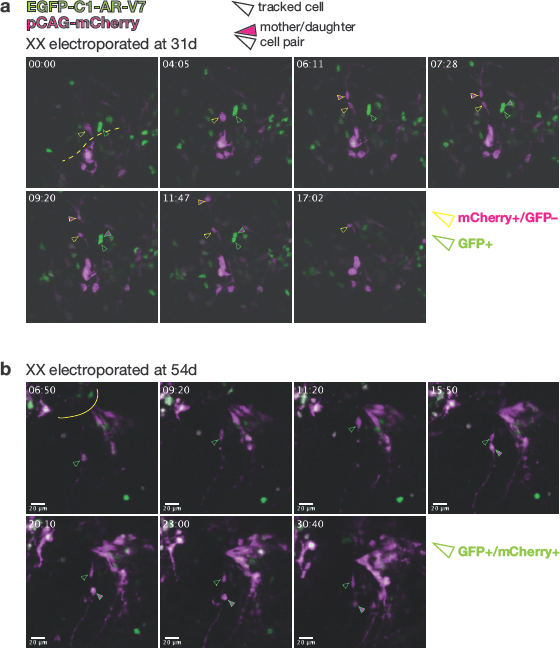
Still images from live imaging of organoids electroporated at a) 31d and b) 54d, and imaged beginning at 2 days post-electroporation (see **Methods** for details of image acquisition). pCAG-mCherry+ cells are shown in magenta, and EGFP-C1-AR-V7+ are shown in green. Green arrowheads: GFP+ cells. Yellow arrowheads: mCherry+/GFP– cells. Filled in arrowheads: one of the tracked daughter cells. a) Comparison of the behaviour of mCherry+ and GFP+ only cells at 33d. Basal surface is up. mCherry+ only cell, upon division (06:11) produces two daughter cells, one of which migrates basally, indicating a more differentiated identity, while the other stays in the ventricular zone (VZ), representing an example of asymmetric division. GFP+ cell divides (07:28), but both of the daughter cells continue to reside in the VZ. b) Behaviour of an mCherry+/GFP+ cell at 56d. Basal surface is down. After division (15:50), both of the daughter cells remain in the VZ. Yellow dashed lines demarcate the apical surfaces. Scale bars: a), b) 20μm. Time scale: hours:minutes.

**Extended Data Figure 5.**
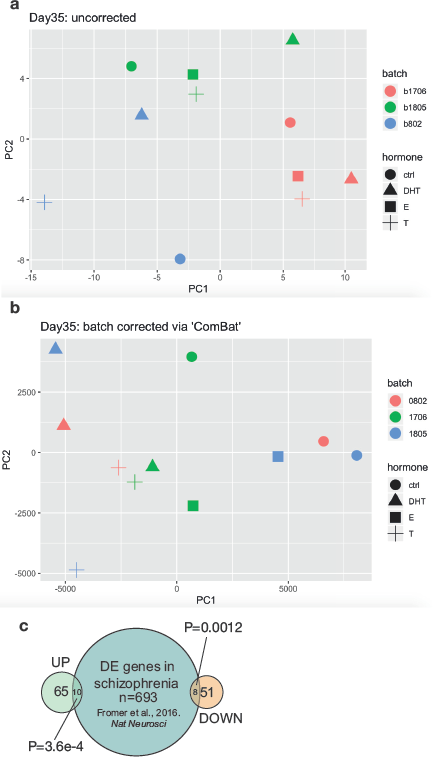
a), b) Principle component analysis (PCA) plots of XX 35d RNA-seq for both a) uncorrected and b) corrected for batch effects. c) Venn diagram representing the overlap of up- and downregulated differentially expressed genes (DEGs) in androgen treated XX 35d organoids with genes differentially expressed in schizophrenia patients^42^. *P* values represent significance of the intersection between the datasets. DE: differentially expressed.

**Extended Data Movie 1.**

Live imaging of XX organoids, electroporated at 31d, and imaged at 33d for >48 hours. Cells labelled with pCAG-mCherry are in magenta. Cells electroporated with EGFP-C1-AR-V7 are in green. Still images from this movie are shown in Extended Data Figure 4a. Time scale: hours:minutes.

**Extended Data Movie 2.**

Live imaging of XX organoids, electroporated at 54d, and imaged at 56d for >48 hours. Cells labelled with pCAG-mCherry are in magenta. Cells electroporated with EGFP-C1-AR-V7 are in green. Still images from this movie are shown in Extended Data Figure 4b. Time scale: hours:minutes.

**Extended Data Movie 3.**

Green channel (EGFP-C1-AR-V7) from Extended Data Movie 2. for easier tracking of electroporated, GFP+ cells. Time scale: hours:minutes.

**Extended Data Table 1.** List of differentially expressed genes between control and androgen-treated organoids Up- and downregulated differentially expressed genes (DEGs) between control and androgen (DHT and T) -treated XX 35d organoids.

| UPREGULATED DEGs |  |  |  |  |
| --- | --- | --- | --- | --- |
| ESEMBL ID | Gene name | Full name | SFARI gene (Gene score ≥3) | DE in SZ (SZDB) |
| ENSG00000065978 | YBX1 | Y-Box Binding Protein 1 |  |  |
| ENSG00000066629 | EML1 | EMAP Like 1 |  |  |
| ENSG00000070785 | EIF2B3 | Eukaryotic Translation Initiation Factor 2B Subunit Gamma |  |  |
| ENSG00000082068 | WDR70 | WD Repeat Domain 70 |  |  |
| ENSG00000089009 | RPL6 | Ribosomal Protein L6 |  |  |
| ENSG00000100029 | PES1 | Pescadillo Ribosomal Biogenesis Factor 1 |  | Yes |
| ENSG00000100281 | HMGXB4 | HMG-Box Containing 4 |  | Yes |
| ENSG00000104738 | MCM4 | Minichromosome Maintenance Complex Component 4 | Yes |  |
| ENSG00000106628 | POLD2 | DNA Polymerase Delta 2, Accessory Subunit |  |  |
| ENSG00000108179 | PPIF | Peptidylprolyl Isomerase F (Cyclophilin D) |  |  |
| ENSG00000109919 | MTCH2 | Mitochondrial Carrier 2 |  | Yes |
| ENSG00000110080 | ST3GAL4 | ST3 Beta-Galactoside Alpha-2,3-Sialyltransferase 4 |  |  |
| ENSG00000116161 | CACYBP | Calcylin Binding Protein |  |  |
| ENSG00000116514 | RNF19B | Ring Finger Protein 19B |  | Yes |
| ENSG00000121152 | NCAPH | Non-SMC Condensin I Complex Subunit H |  |  |
| ENSG00000121390 | PSPC1 | Paraspeckle Component 1 |  |  |
| ENSG00000121680 | PEX16 | Peroxisomal Biogenesis Factor 16 |  | Yes |
| ENSG00000123395 | ATG101 | Autophagy Related 101 |  |  |
| ENSG00000123416 | TUBA1B | Tubulin Alpha 1b |  |  |
| ENSG00000124688 | MAD2L1BP | MAD2L1 Binding Protein |  |  |
| ENSG00000129680 | MAP7D3 | MAP7 Domain Containing 3 |  |  |
| ENSG00000130021 | PUDP | Pseudouridine 5'-Phosphatase |  |  |
| ENSG00000132341 | RAN | RAN, Member RAS Oncogene Family |  |  |
| ENSG00000134057 | CCNB1 | Cyclin B1 |  |  |
| ENSG00000137691 | C11orf70 (CFAP300) | Cilia And Flagella Associated Protein 300 |  |  |
| ENSG00000137876 | RSL24D1 | Ribosomal L24 Domain Containing 1 |  |  |
| ENSG00000138231 | DBR1 | Debranching RNA Lariats 1 |  |  |
| ENSG00000138764 | CCNG2 | Cyclin G2 |  |  |
| ENSG00000139921 | TMX1 | Thioredoxin Related Transmembrane Protein 1 |  |  |
| ENSG00000143575 | HAX1 | HCLS1 Associated Protein X-1 |  |  |
| ENSG00000145545 | SRD5A1 | Steroid 5 Alpha-Reductase 1 |  | Yes |
| ENSG00000148229 | POLE3 | DNA Polymerase Epsilon 3, Accessory Subunit |  | Yes |
| ENSG00000152465 | NMT2 | N-Myristoyltransferase 2 |  |  |
| ENSG00000156508 | EEF1A1 | Eukaryotic Translation Elongation Factor 1 Alpha 1 |  |  |
| ENSG00000162704 | ARPC5 | Actin Related Protein 2/3 Complex Subunit 5 |  |  |
| ENSG00000164087 | POC1A | POC1 Centriolar Protein A |  |  |
| ENSG00000164109 | MAD2L1 | Mitotic Arrest Deficient 2 Like 1 |  |  |
| ENSG00000164758 | MED30 | Mediator Complex Subunit 30 |  |  |
| ENSG00000165526 | RPUSD4 | RNA Pseudouridine Synthase D4 |  |  |
| ENSG00000165775 | FUNDC2 | FUN14 Domain Containing 2 |  |  |
| ENSG00000166347 | CYB5A | Cytochrome B5 Type A |  |  |
| ENSG00000166557 | TMED3 | Transmembrane P24 Trafficking Protein 1 |  |  |
| ENSG00000166851 | PLK1 | Polo Like Kinase 1 |  |  |
| ENSG00000167085 | PHB | Prohibitin | Yes |  |
| ENSG00000167112 | TRUB2 | TruB Pseudouridine Synthase Family Member 2 |  |  |
| ENSG00000168282 | MGAT2 | Alpha-1,6-Mannosyl-Glycoprotein 2-Beta-N-Acetylglucosaminyltransferase |  |  |
| ENSG00000169714 | CNBP | CCHC-Type Zinc Finger Nucleic Acid Binding Protein |  |  |
| ENSG00000170364 | SETMAR | SET Domain And Mariner Transposase Fusion Gene |  |  |
| ENSG00000171720 | HDAC3 | Histone Deacetylase 3 |  |  |
| ENSG00000172053 | QARS | Glutamyl-tRNA Synthetase |  | Yes |
| ENSG00000176903 | PNMA1 | PNMA Family Member 1 |  |  |
| ENSG00000180581 | SRP9P1 | Signal Recognition Particle 9 Pseudogene 1 |  |  |
| ENSG00000182054 | IDH2 | Isocitrate Dehydrogenase (NADP(+)) 2 |  | Yes |
| ENSG00000183431 | SF3A3 | Splicing Factor 3a Subunit 3 |  |  |
| ENSG00000183684 | ALYREF | Aly/REF Export Factor |  |  |
| ENSG00000186665 | C17orf58 | Chromosome 17 Open Reading Frame 58 |  |  |
| ENSG00000196262 | PPIA | Peptidylprolyl Isomerase A |  |  |
| ENSG00000196591 | HDAC2 | Histone Deacetylase 2 |  | Yes |
| ENSG00000198039 | ZNF273 | Zinc Finger Protein 273 |  |  |
| ENSG00000229119 | AC026403.1 | 60S acidic ribosomal protein (RPLP0) pseudogene |  |  |
| ENSG00000244563 | RPS26P19 | Ribosomal protein S26 pseudogene 19 |  |  |
| ENSG00000251668 | AC113404.2 | pseudogene similar to part of microtubule associated serine/threonine kinase 2 MAST2 |  |  |
| ENSG00000255302 | EID1 | EP300 Interacting Inhibitor Of Differentiation 1 |  |  |
| ENSG00000270800 | RPS10-NUDT3 | RPS10-NUDT3 readthrough |  |  |
| ENSG00000280071 | GATD3B | Glutamine amidotransferase like class 1 domain containing 3B |  |  |

| DOWNREGULATED DEGs |  |  |  |  |
| --- | --- | --- | --- | --- |
| ESEMBL ID | Gene name | Full name | SFARI gene (Gene score ≥3) | DE in SZ (SZDB) |
| ENSG0000015568 | RGPD5 | RANBP2 Like And GRIP Domain Containing 5 |  | Yes |
| ENSG00000092929 | UNC13D | Unc-13 Homolog D |  |  |
| ENSG00000099840 | IZUMO4 | IZUMO Family Member 4 |  |  |
| ENSG00000100596 | SPTLC2 | Serine Palmitoyltransferase Long Chain Base Subunit 2 |  |  |
| ENSG00000101104 | PABPC1L | Poly(A) Binding Protein Cytoplasmic 1 Like |  |  |
| ENSG00000107317 | PTGDS | Prostaglandin D2 Synthase |  | Yes |
| ENSG00000108515 | ENO3 | Enolase 3 |  |  |
| ENSG00000112855 | HARS2 | Histidyl-TRNA Synthetase 2, Mitochondrial |  |  |
| ENSG00000116266 | STXBP3 | Syntaxin Binding Protein 3 |  |  |
| ENSG00000123473 | STIL | STIL Centriolar Assembly Protein |  |  |
| ENSG00000129003 | VPS13C (PARK23) | Vacuolar Protein Sorting 13 Homolog C |  |  |
| ENSG00000132485 | ZRANB2 | Zinc Finger RANBP2-Type Containing 2 |  | Yes |
| ENSG00000136100 | VPS36 | Vacuolar Protein Sorting 36 Homolog |  |  |
| ENSG00000137831 | UACA | Uveal Autoantigen With Coiled-Coil Domains And Ankyrin Repeats |  |  |
| ENSG00000139219 | COL2A1 | Collagen Type II Alpha 1 Chain |  |  |
| ENSG00000143416 | SELENBP1 | Selenium Binding Protein 1 |  | Yes |
| ENSG00000147654 | EBAG9 | Estrogen Receptor Binding Site Associated Antigen 9 |  |  |
| ENSG00000148600 | CDHR1 | Cadherin Related Family Member 1 |  |  |
| ENSG00000154222 | CC2D1B | Coiled-Coil And C2 Domain Containing 1B |  |  |
| ENSG00000157036 | EXOG | Exo/Endonuclease G |  |  |
| ENSG00000157379 | DHRS1 | Dehydrogenase/Reductase 1 |  |  |
| ENSG00000160194 | NDUFV3 | NADH:Ubiquinone Oxidoreductase Subunit V3 |  | Yes |
| ENSG00000161298 | ZNF382 | Zinc Finger Protein 382 |  |  |
| ENSG00000167264 | DUS2 | Dihydrouridine Synthase 2 |  |  |
| ENSG00000168300 | PCMTD1 | Protein-L-Isoaspartate (D-Aspartate) O-Methyltransferase Domain Containing 1 |  |  |
| ENSG00000176809 | LRRC37A3 | Leucine Rich Repeat Containing 37 Member A3 |  |  |
| ENSG00000179889 | PDXDC1 | Pyridoxal Dependent Decarboxylase Domain Containing 1 |  |  |
| ENSG00000181085 | MAPK15 | Mitogen-Activated Protein Kinase 15 |  |  |
| ENSG00000181666 | HKR1 (ZNF875) | Zinc Finger Protein 875 |  |  |
| ENSG00000188234 | AGAP4 | ArfGAP With GTPase Domain, Ankyrin Repeat And PH Domain 4 |  |  |
| ENSG00000196275 | GTF2IRD2 | GTF2I Repeat Domain Containing 2 |  |  |
| ENSG00000196739 | COL27A1 | Collagen Type XXVII Alpha 1 Chain |  |  |
| ENSG00000198804 | MT-CO1 | Mitochondrially Encoded Cytochrome C Oxidase I |  |  |
| ENSG00000198840 | MT-ND3 | Mitochondrially Encoded NADH:Ubiquinone Oxidoreductase Core Subunit 3 |  |  |
| ENSG00000198888 | MT-ND1 | Mitochondrially Encoded NADH:Ubiquinone Oxidoreductase Core Subunit 1 |  | Yes |
| ENSG00000204681 | GABBR1 | Gamma-Aminobutyric Acid Type B Receptor Subunit 1 |  |  |
| ENSG00000205583 | STAG3L1 | Stromal Antigen 3-Like 1 (Pseudogene) |  |  |
| ENSG00000225630 | MTND2P28 | MT-ND2 Pseudogene 28 |  |  |
| ENSG00000226232 | NPIPB14P | Nuclear pore complex interacting protein family member B14, pseudogene |  |  |
| ENSG00000228253 | MT-ATP8 | Mitochondrially encoded ATP synthase membrane subunit 8 |  |  |
| ENSG00000232196 | MTRNR2L4 | MT-RNR2 Like 4 |  |  |
| ENSG00000234441 | AC105272.1 |  |  |  |
| ENSG00000237732 | CT75 | Cancer/testis associated transcript 75 |  |  |
| ENSG00000237973 | MTCO1P12 | MT-CO1 pseudogene 12 |  |  |
| ENSG00000239912 | RPL39P36 | Ribosomal protein L39 pseudogene 36 |  |  |
| ENSG00000243716 | NPIPB5 | Nuclear pore complex interacting protein family member B5 |  |  |
| ENSG00000248527 | MTATP6P1 | MT-ATP6 pseudogene 1 |  |  |
| ENSG00000256618 | MTRNR2L1 | MT-RNR2 Like 1 |  |  |
| ENSG00000258366 | RTEL1 | Regulator Of Telomere Elongation Helicase 1 |  |  |
| ENSG00000258890 | CEP95 | Centrosomal Protein 95 |  |  |
| ENSG00000274615 | AC233968.1 | Aminopeptidase puromycin sensitive (NPEPPS) pseudogene |  |  |

